# Soft biocompatible polymer optical fiber tapers for implantable neural devices

**DOI:** 10.1101/2024.10.29.620629

**Authors:** Marcello Meneghetti, Jiachen Wang, Kunyang Sui, Rune W. Berg, Christos Markos

## Abstract

Optical fibers are between the most common implantable devices for delivering light in the nervous system for optogenetics and infrared neural stimulation applications. Tapered optical fibers, in particular, can offer homogeneous light delivery to a large volume and spatially resolved illumination compared to standard flat-cleaved fibers while being minimally invasive. However, the use of tapers for neural applications has up to now been limited to silica optical fibers, whose large Young’s modulus can cause detrimental foreign body response in chronic settings. Here, we present the fabrication and optimization of tapered fiber implants based on polymer optical fibers (POFs). After numerically determining the optimal materials and taper geometry, we fabricated two types of POFs by thermal fiber drawing. The fabrication of the taper was achieved by chemical etching of the fibers, for which several solvents previously reported in literature have been tested. The influence of different parameters on the etching process and on the quality of the obtained tapers was also investigated. The large illumination volume of the produced high-quality taper-based implants was finally tested *in vitro* in a brain phantom.

## Introduction

Since the early 2000s, with the demonstration of optogenetics and infrared neurostimulation (INS) (1–3), optical methods to modulate neuronal activity have become increasingly popular. This created the need for ways to efficiently deliver light in different brain region, with optical fibers emerging as one of the most promising solutions (4, 5). Tapered optical fibers, or tapers, have been recently attracting interest for this application. Unlike standard flat-cleaved optical fibers, which are only able to illuminate brain regions in close proximity to their end facets, tapers exhibit a large illumination volume along their sharp profile(6). Furthermore, the modal propagation properties of light in tapered profiles allow for depth-selective illumination by modifying the input coupling, while being minimally invasive thanks to their smaller average cross-section (6, 7). The optical fiber tapers reported up to now for neural applications, however, are based on silica glass, which has a Young’s modulus orders of magnitudes larger than the one of brain tissue (8). This large mismatch in mechanical properties represents a challenge in the use of silica and other stiff materials as the main constituent for brain implants, due to the arisal of inflammation, glial scarring and strong foreign body response in chronic settings (9).

As a way to mitigate this issue, a great deal of research effort has been devoted to the development of biocompatible neural implants based on soft materials such as polymers and hydrogels (10–12). Polymer optical fibers (POFs), in particular, thanks to their small footprint and low bending stiffness, have been demonstrated to be an effective platform for both optogenetics and INS (13–16). Moreover, thanks to the versatility of POF fabrication by thermal drawing process and post-processing by various micro- and nano-fabrication techniques, several functionalities such as stimulation, recording, drug delivery and sensing can be monolithically integrated in POF-based implants and tapers (17–23).

In this work, we designed and fabricated POF-based tapers optimized for efficient light delivery in brain tissue. Polycarbonate (PC) and poly(methyl methacrylate) (PMMA) have been chosen as starting materials due to their high transmittance in the visible part of the spectrum, which makes them compatible with optogenetics applications (24, 25). Chemical etching was then used to create a taper profile at the tip of the in-house fabricated POFs (Fig. 1a) in order to achieve the desired geometry, which had been previously numerically determined by finite element method (FEM) simulations. Several solvents have been tested, and the etching parameters have been optimized to eliminate different kinds of defects arising from the chemical etching process. We identified that the most suitable solvents that allowed us to repeatably fabricate high quality POF-based tapers suitable for optogenetic modulation of neurons in large brain structures *in vivo*.

**Fig 1.**
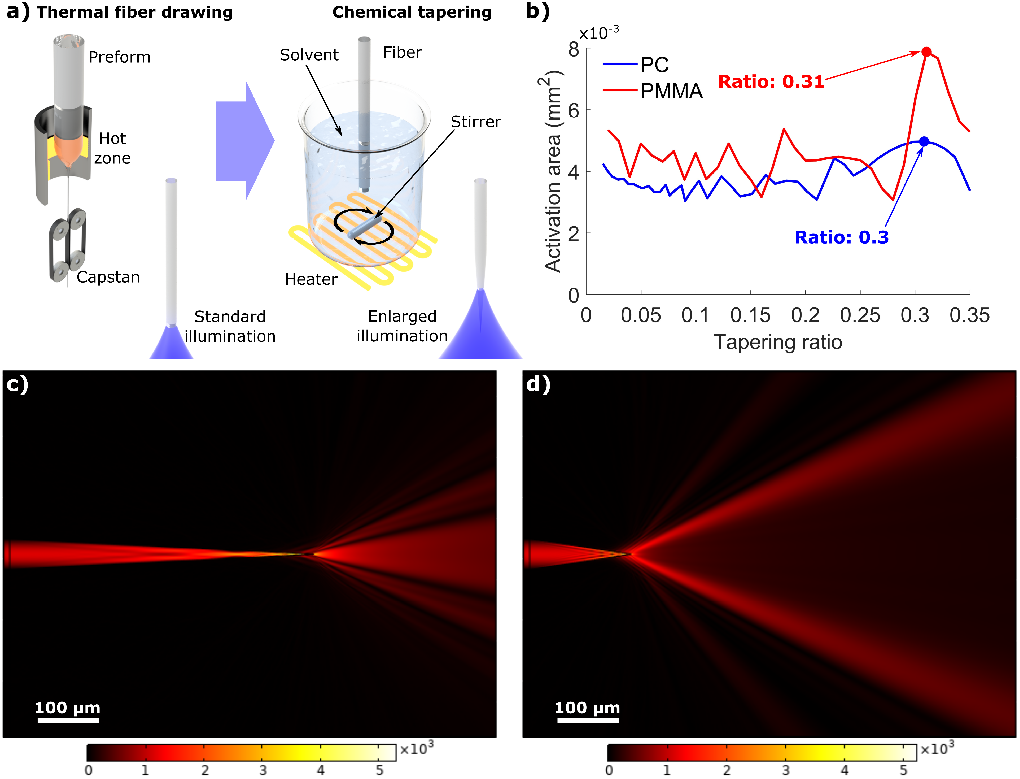
a) Schematics of the process: a single index POF produced by thermal fiber drawing is chemically tapered to enlarge the illumination area; b) Results of the numerical optimization process, showing that for both PC and PMMA a taper ratio of 0.3 maximizes the activation region; c,d) Simulated electric field distribution for the light delivered to the brain from tapers with taper ratios of 0.1 and 0.3, respectively.

## Results and Discussion

As a starting point for the development of the taper, we used FEM simulations to determine the ideal geometry for achieving minimal inflammatory response from the tissue while ensuring efficient large area illumination of the brain. Based on the mechanical simulation model developed by our group and previously reported in (26), we found a fiber diameter of 50 *µm* at the implant’s largest point to have optimal bending stiffness while still allowing for implantation without buckling (12). Once the initial diameter had been fixed, we proceeded to define the appropriate tapering ratio *R*_*t*_ (ratio between the diameter at the largest point and the length of the tapered section). A radiant exposure of 1000 *W/m*^2^ was fixed as the minimum threshold for the optogenetic modulation of neuronal activity, and a FEM model was then used to determine which tapering ratio corresponded to the largest effective volume of tissue reaching the threshold. We thus determined the ideal ratio to be of 0.30 and 0.31 for PC and PMMA, respectively (Fig. 1b). The presence of a definite optimal ratio is compatible with the simulated light at the taper output: an increased ratio corresponds to a broader illumination angle, but lower average intensity (Fig.1c,d).

In order to identify the best solvents to be used for tapering our fibers, we tested a number of different solvents that have been used in literature to etch or dissolve polycarbonate and PMMA (27–29). Before immersing the fibers in the solvent, a critical step was to anneal them under vacuum for 24 hours at 120°C (PC) and 90°C (PMMA) to release internal stress, which we observed to cause cracking during etching. Scanning electron microscopy (SEM) images of typical examples of a non- and a pre-annealed fibers that were etched using the same conditions is shown in Fig. 2a,b. Despite the annealing, however, many of the tested solvents caused damage to the fibers in the form of swelling (Fig. 2c), increased roughness (Fig. 2d), loss of transparency in the etched part (Fig. 2e) or destruction of the fiber’s surface (Fig. 2f).

**Fig 2.**
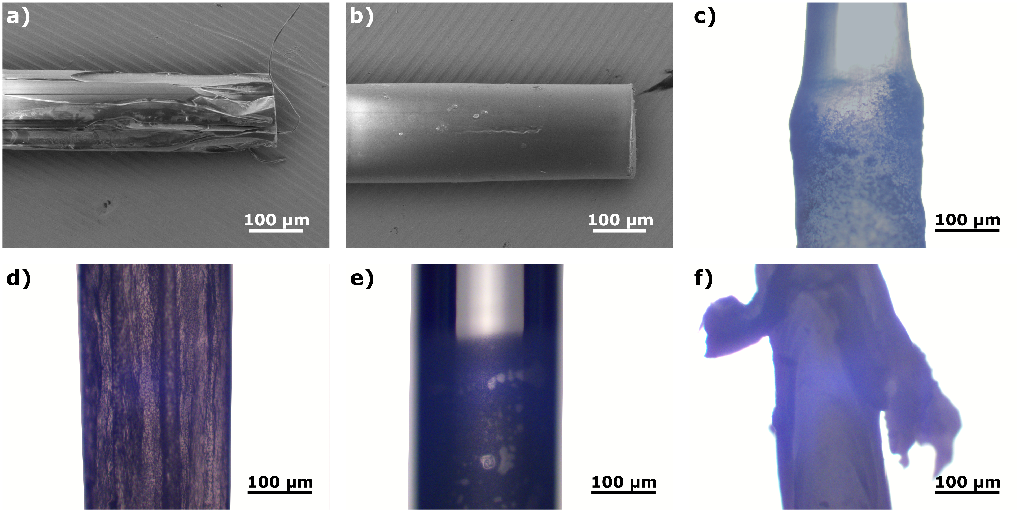
a,b) SEM images of a PMMA fiber etched in *ace* prior to and after annealing under vacuum, respectively; c-f) Damage caused by etching polycarbonate fibers in *ace, chlo, dcmet* and dimethylformamide, respectively.

After extensive evaluation of the damage caused for each combination of solvent and polymer, we found cyclohexanone (*chex*) to be a suitable solvent for PC. At this stage, the solvents found to be viable for testing with PMMA were 1,2- dichlorobenzene (*dcben*), acetone (*ace*), chloroform (*chlo*), dibromomethane (*dbmet*), dichloromethane (*dcmet*), n-butyl acetate (*nbace*), toluene (*tol*) and trichloroethylene (*tceth*). To controllably etch the fibers into tapers with the desired geometry, a crucial step is the determination of the etching rate *v*_*e*_. This parameter was investigated by a “step-etching” method, in which a large diameter fiber was initially immersed in the solvent and then retracted by 500 *µm* at regular time intervals, thus creating a step profile (Fig. 3a, inset). Here, the retraction was performed by using a precision 3-axis translational stage (Thorlabs MBT616D/M, resolution of 50 *µm/rev*) while the solvent was kept in motion by a magnetic stirrer. All of the *v*_*e*_ measurements were performed at room temperature and with a stirring speed of 200 *rpm*.

**Fig 3.**
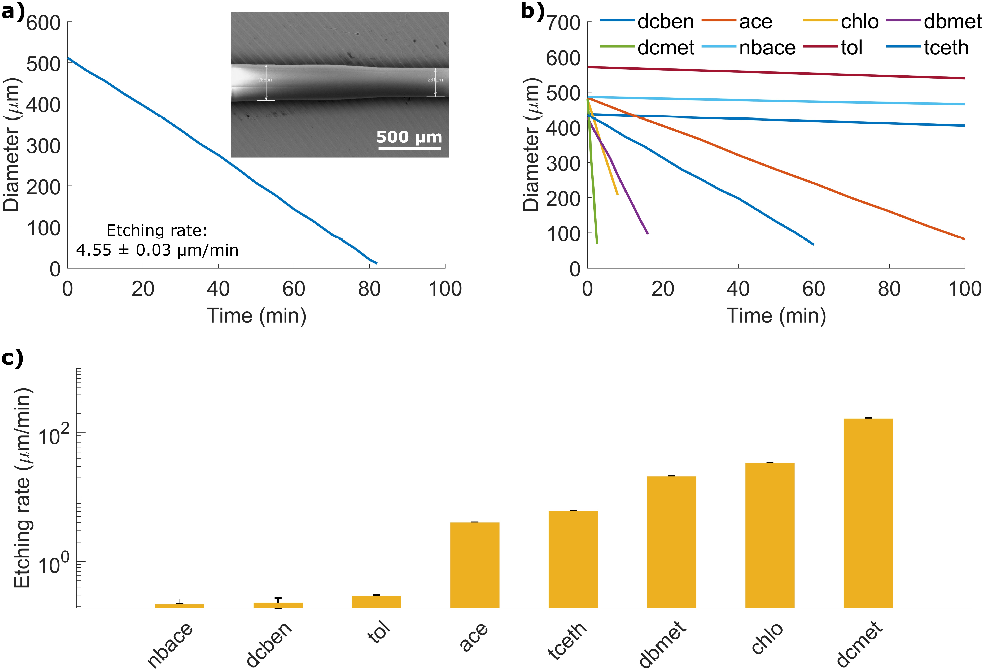
a) Diameter variation of a polycarbonate fiber at different immersion times in *chex* (inset: SEM image of one of the fibers used for the evaluation of the etching rate); b) Diameter variation of PMMA fibers at different immersion times in several solvents; c) Estimated etching rates for the different solvents tested on PMMA.

The diameters corresponding to the different immersion times were then measured by SEM, and used to calculate by linear regression the etching rate of the above mentioned solvents. For *chex*, the solvent chosen for the fabrication of PC tapers, we found the etching rate to be of 4.55*µm/min* (Fig. 3a). A very broad spread of etching rates was found between the solvents tested for PMMA, with the highest (for *dcmet*) being three orders of magnitude faster than the lowest (for *nbace*) (Fig 3b,c). The etching rates for each solvent were used to calculate the retraction speed *v*_*r*_ required to obtain the desired geometry by continuously retracting a fiber immersed in the solvent:

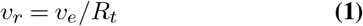

The retraction speed for obtaining a PC taper by etching a fiber in *chex* was thus found to be 15.17 *µm/min*, while the values of *v*_*r*_ for the PMMA taper fabrication vary from 0.70 *µm/min* to 535 *µm/min*, depending on the solvent. While choosing a solvent, one must consider that an exceedingly slow retraction speed would render the tapering process unpractical in terms of time consumption, while a too fast etching could negatively impact the stability of the process. Thus, *ace* and *tceth* were initially considered as candidates for the fabrication of the PMMA taper. However, we repeatedly observed the formation of large material aggregates along the etched length of the fiber (Fig. 4a). In an attempt to solve this issue, which we attributed to a too long permanence of the fiber in the solvent at each step, we proceeded to tune the etching speed of the solvents. This can be done either by changing the solvent temperature or the stirring speed. The relationships between these parameters and the etching speeds, measured by step-etching PMMA in *ace*, are presented in Fig. 4b. A second hypothesis for the formation of aggregates was a misalignment of the fiber with respect to the center of the beaker under stirring. This would result in a perturbation of the fluid motion, creating a region of slow flow on one side of the fiber (Fig. 4c and Supplementary Video 1), which, according to the FEM simulations we performed, would expose one side of the fiber to speeds ∼ 20 times lower than the opposite side (Fig. 4d).

**Fig 4.**
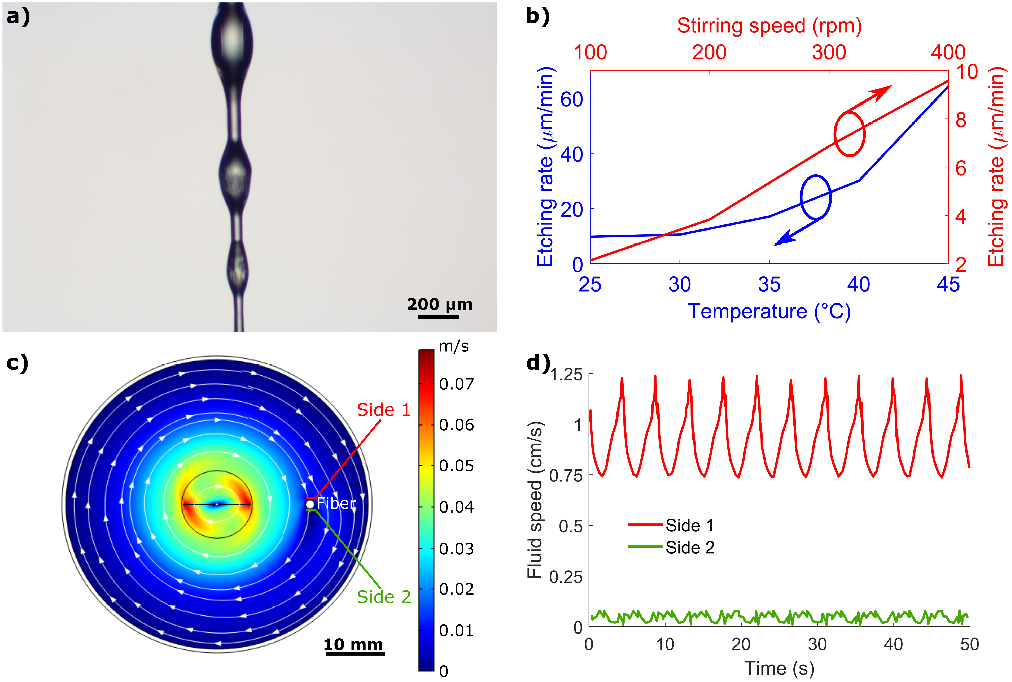
a) Optical microscope image of aggregates forming on a PMMA fiber etched in *tceth*; b) Relationship between solvent temperature and etching rate (blue line and axes) and between stirring speed and etching rate (red line and axes); c) Simulation of the flow of solvent perturbed by an off-center fiber during stirring d) Difference in the solvent flow speed experienced by two opposite sides of an off-center fiber.

Thus, the alignment between the fiber and the beaker was optimized to ensure an isotropic etching. None of these approaches, however, resulted in repeatable, high quality PMMA taper fabrication. The desired geometry was finally obtained by changing the solvent to *dbmet*, thus confirming that efficient tapering of PMMA POFs requires higher etching speeds than PC.

By using the previously described process, we were able to obtain high quality tapers from both PC and PMMA, with measured taper ratios of 0.302 and 0.307, respectively, very close to the desired values of 0.30 and 0.31 (Fig. 5a,b). To verify the expected large illumination area, the tapers were immersed in thin slices of brain phantom (0.6% w/v agarose in water), a material known to have optical properties similar to the ones of brain tissue (8). Light from a supercontinuum source (NKT SuperK Extreme, 100 mW output power) was coupled into the tapers through a silica patch cable, and illumination maps were recorded using a digital microscope (Dino-Lite, AM7915MZT) using an experimental setup configuration similar to the one described in (8). Both tapers exhibit a large illumination angle, with a slightly asymmetric profile that can be attributed to either variations in the coupling from the patch cable or to anisotropies in the brain phantom slice (Fig. 5c,d).

**Fig 5.**
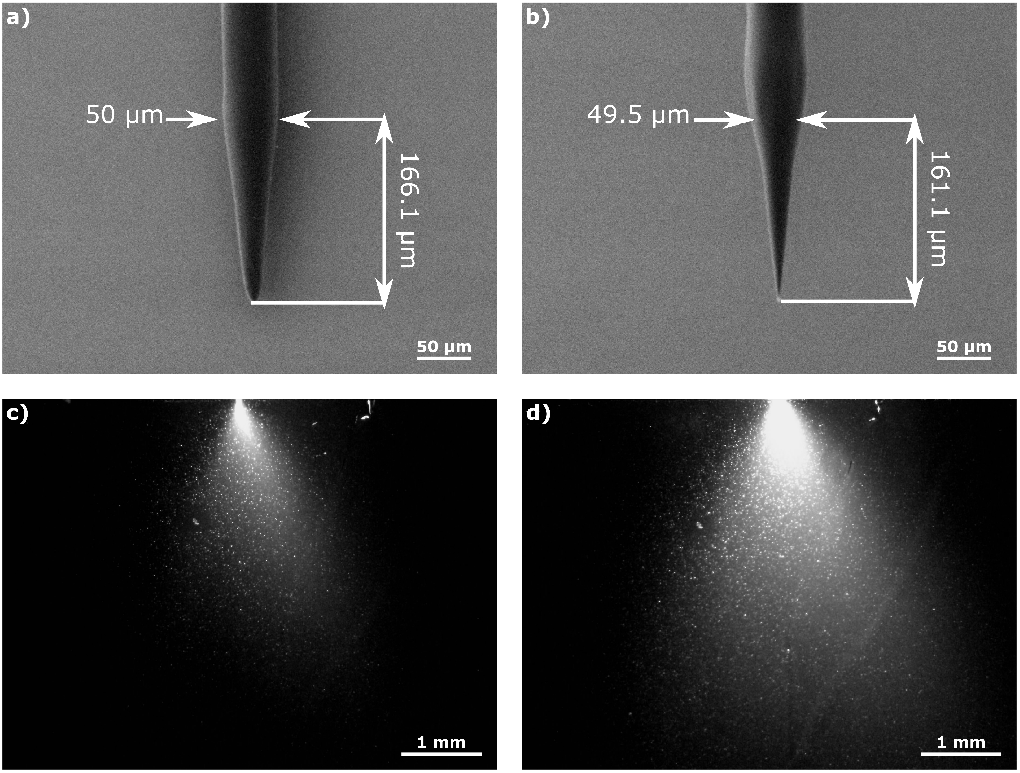
a,b) SEM images of the PC and PMMA tapers, respectively; c,d) Illumination map of the white light delivered in brain phantom by the tapered tip of PC and PMMA fibers, respectively.

To better analyze the effective illumination profile, the recorded illumination maps were normalized to their maximum intensity and plotted in the form of isocontours depicting the decay of intensity in space down to 10% of its maximum.

For the PC fiber, we observed a maximum lateral spread of ∼1.6 mm and a penetration depth of ∼2.2 mm at a 10% level (Fig. 6a). By observing the 1-dimensional intensity profiles along orthogonal lines intersecting at the maximum intensity points, we estimated the propagation decay in brain phantom to be on average 28.3 dB/mm laterally and 5.4 dB/mm along the taper’s axis (Fig. 6b,c). In comparison, light from the PMMA taper spread ∼2.6 mm laterally and∼ 3.3 mm longitudinally (Fig. 6d). This larger illumination volume is compatible with the lower measured decay rates (average of 13.9 dB/mm lateral, 3.5 dB/mm longitudinal, as shown in Fig. 6e,f) and thus doesn’t appear to be caused by a larger amount of light being coupled in the PMMA taper, but rather by the lower confinement of light within the taper caused by the lower refractive index contrast between PMMA and the surrounding brain phantom.

**Fig 6.**
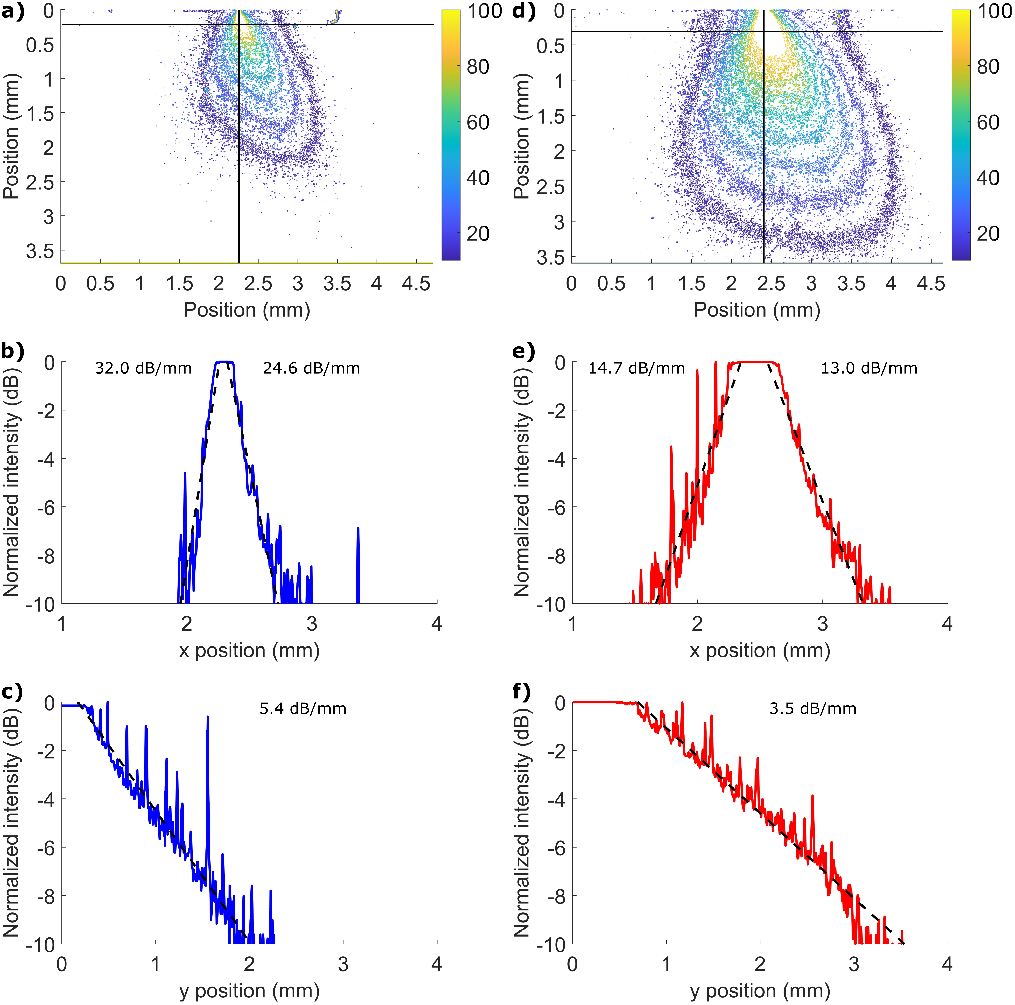
a) Isocontours showing decay of the light delivered by a PC taper, extracted from Fig.2c; b,c) Respectively, transverse and longitudinal profiles (horizontal and vertical cut lines in (a)) of the light delivered by a PC taper, normalized and converted to decibels, with linear fits of the light decay (black dashed lines); d) Isocontours showing decay of the light delivered by a PMMA taper, extracted from Fig.2d; e,f) Respectively, transverse and longitudinal profiles (horizontal and vertical cut lines in (d)) of the light delivered by a PMMA taper, normalized and converted to decibels, with linear fits of the light decay (black dashed lines).

## Conclusions

In summary, in this work we have investigated the use of chemical etching to develop tapered polymer optical fibers designed to achieve illumination of large populations of neurons while minimizing the foreign body response caused by the mechanical mismatch between brain tissue and implants. During the optimization of the etching process, we found that most common solvents used for etching PC have a negative impact on the quality of the optical fibers, whereas the diameter of PMMA POFs can be reduced using a broad range of chemicals with significantly varying etching rates. This flexibility of processing proved to be an important advantage, since we show here that the possibility of controlling the speed of the chemical etching, either by selecting the proper solvent or by fine-tuning the temperature and mixing speed during the process, is crucial for achieving high quality tapers. The process developed during this investigation allowed us to employ *chex* and *dbmet* (for PC and PMMA, respectively) to reliably obtain tapers matching the desired taper ratios, which were determined previously by optimizing the geometry of the implants with FEM optical and mechanical simulations, with deviations of less than 1%. The *in vitro* mapping of the illumination area achievable using the developed tapers, performed in an agarose gel medium matching the characteristics of the brain, showed that the lower refractive index PMMA polymer can illuminate a larger brain region when compared with PC, with an estimated penetration depth of more than 3 mm at −10 dB.

In conclusion, we believe that the soft polymeric tapers presented in this work have high potential as implantable devices for optical neuromodulation and fiber photometry in large neuronal population, especially in chronic settings where standard silica-based implants suffer from high levels of gliosis and foreign body response.

## ACKNOWLEDGEMENTS

This research has been financially supported by Lundbeck Fonden projects Multi-BRAIN (R276-2018-869) and R380-2021-1171 and EIC Pathfinder Open Project 101130161 - Move2Treat.

We would like to express our gratitude to Jakob Janting (DTU Electro) for his support during this investigation.

## DATA AVAILABILITY

Data underlying the results presented in this paper are not publicly available at this time but may be obtained from the authors upon reasonable request.

## Supplementary Video 1: Perturbation in the solvent’s flow speed during stirring

Graphical representation of the solvent’s flow speed during stirring when the polymer optical fiber (black dot) is at the center of the beaker. The color scalebar is the same as Fig. 4c.

